# Spectral Prompting: Unsupervised Recovery of Human Hair Follicle Cell-Type and Multiscale Systems Architecture from Bulk and Single-Cell RNA-Seq Datasets via Single-Gene Seeded Spectral Unfolding

**DOI:** 10.64898/2026.05.19.726151

**Authors:** Talveen S. Purba

## Abstract

Bulk RNA sequencing datasets are assumed to carry minimal resolvable programmatic and cell type biological information; as such, in the absence of single-cell resolution, researchers prioritise data analysis approaches based on differential expression, or rely on deconvolution and co-expression methods that require external reference panels, large multi-sample cohorts, or prior single-cell data to resolve cell-type structure. Here I describe the recovery of specialised cell-type and systems gene expression architecture resolved from a static gene expression dataset of untreated cultured human hair follicles (pooled from N=12 patients) isolated from scalp skin. To achieve this, I used graph theoretic methods to mathematically transform gene expression data into a latent space of relational structure, which was spectrally organised into coarse- and fine-grained modes and partitioned using a purpose-built computational algorithm. This permitted the synthesis of a computational Spectral Prompting system, whereby a single gene can be seeded to “unfold” to reveal associated partners across manifold projections in gene expression space. Individual projections across the manifold can reveal rich individual gene expression programmes, which can then be aggregated to identify core-associated genes for a given spectral gene prompt, both within the manifold analysed and across *>*1 manifold constructions. With this, I recover hitherto unresolved gene expression programmes from bulk data, including, but not limited to, epithelial hair follicle stem cell (eHFSC), hair shaft, dermal papilla and endothelial gene expression signatures. Focusing on querying KRT15, a human anagen bulge eHFSC and progenitor marker, raw output from individual spectral prompts during testing recovered known eHFSC-associated genes including LGR5, LHX2 and CXCL14, and discovered new candidate human eHFSC and progenitor cell-associated markers, such as RGMA and MUCL1 which were validated *in situ*. Finally, I show a brief demonstration that the technique can be similarly applied to single-cell data (GSE129611), whereby a KRT15 gene prompt from a combined expression matrix was mapped to a KRT15+/CXCL14+/LHX2+/DIO2+/SFRP1+ cell population (31/6000 cells) independent of standard clustering tools. Moving forward, from this foundation, the method will be developed to study how latent gene expression space shifts following perturbation or pathology.

## Background

Current computational advances have centred on foundational transformer models to develop learned insights on a given corpus. These models typically require large amounts of training data, computation and skill to build neural networks and other learned representations. In medicine and the biosciences, such representations can be used in general medical artificial intelligence [1], biomolecule structure prediction [2], and multiomics [3] and cell biology [3–6].

In this preprint I describe Spectral Prompting, a system that uses single gene queries (where the query must be expressed in the target dataset) on a constructed manifold, to “unfold” gene expression programmes in the latent mathematical space. This is achieved without training, supervision, external reference panels, curated databases, or literature-derived annotations, and instead draws upon principles from spectral graph theory [7] to construct and query a gene expression manifold from bulk and single cell RNA-Seq data that can be custom built from a relatively small number of samples (lowest N tested = 4 samples). I refer to this broader computational framework as Resonance Intelligence (RI), and its current implementation as a queryable discovery instrument, Aethereos Biology.

To demonstrate this, I primarily apply Spectral Prompting to human hair follicle biology [8, 9] and human epithelial stem cell biology [10, 11], constructing a graph from a pooled dataset of control untreated hair follicle samples from our published [12] and unpublished bulk datasets, alongside a demonstration using single cell human hair follicle data, revealing how Spectral Prompting recovers coherent biological gene expression programmes in hair biology.

## Methods

### Human Hair Follicle Culture, RNA-Seq and immunofluorescence

Human scalp hair follicles were dissected from scalp tissue donated by consenting patients undergoing hair restoration procedures. Samples were handled in compliance with HTA regulations under University of Manchester and NHS REC ethical approval. Samples were either cultured untreated for 48h and processed for RNA-Seq, or isolated following dissection, frozen and processed for immunofluorescence staining as described previously [12]. Frozen human hair follicle tissue sections were stained using antibodies against CXCL14 (Abcam, ab264467), MUCL1 (SBEM) (Novus, NBP1-92366) and RGMA (Invitrogen, MA5-57221).

### Spectral Graph Construction of the Gene Manifold and Spectral Prompting

Gene expression read count data from bulk RNA-Seq (and Raw UMI counts for the Single-Cell example using GSE129611) were mathematically transformed to encode gene-gene relational structure across samples. From this, a weighted nearest-neighbour graph was constructed over the gene set and its graph Laplacian spectrally decomposed, embedding all genes into a continuous latent space. The eigenvalue spectrum was partitioned into levels from coarse to fine resolution modes.

Spectral Prompting first seeds a single expressed gene of interest (GOI). A recursive refinement algorithm descends from coarse to fine resolution, reconstructing the manifold at multiple steps, yielding a neighbourhood for a given GOI defined by pre-set exclusionary criteria for mode membership.

## Results & Discussion

### Spectral Prompting Recovers Known Hair Follicle and Cell Biology Paradigms From Bulk Data

To demonstrate the Spectral Prompting system, I arbitrarily selected markers relevant to human hair follicle biology, namely hair shaft cortex differentiation (KRT85) [13] and associated signalling (LEF1) [14], epithelial stem cells (KRT15) [10], the hair follicle mesenchyme (VCAN) [15], pigmentation (DCT, PMEL) [16], cell cycle (MKI67) [17], hair follicle vascular biology (PECAM1) [18], and finally MPZL3, recently described to localise to the inner root sheath [9].

Next, I randomly selected a spectral level (within a constructed manifold by uniform random draw) where the GOI was present, and seeded the query GOI and executed the designed algorithm without pre-selection of associated genes/pathways, or without filtering (however for interpretation, it is advised that this is filtered by a threshold minimum mean read count) or aggregation (mentioned below). The results of the single raw gene prompts, which comprise several transcripts per query, are demonstrated in Tables 1–9. 7 of the 9 prompts successfully resolved, whereas 2 (MKI67 and PMEL) were degenerate, and resolved on the second random draw.

**Table 1:** Randomised KRT85 spectral prompt. Output from a single randomly drawn spectral level. Expression shown as mean normalised read count.x.

| # | Gene | Expression |
| --- | --- | --- |
| 1 | **KRT85** | 160,365 |
| 2 | KRT35 | 33,429 |
| 3 | LAP3 | 8,181 |
| 4 | PPP2R1B | 5,955 |
| 5 | SP6 | 3,999 |
| 6 | FRMPD1 | 2,262 |
| 7 | LEF1 | 1,848 |
| 8 | DNAJC6 | 1,625 |
| 9 | SLC9A2 | 702 |
| 10 | SGK3 | 363 |
| 11 | PCDH19 | 360 |
| 12 | NKD1 | 333 |
| 13 | NALCN | 108 |

**Table 2:** Randomised LEF1 spectral prompt. Output from a single randomly drawn spectral level. Expression shown as mean normalised read count.

| # | Gene | Expression |
| --- | --- | --- |
| 1 | KRT85 | 160,365 |
| 2 | DSG4 | 8,503 |
| 3 | LAP3 | 8,181 |
| 4 | KRT39 | 6,349 |
| 5 | SP6 | 3,999 |
| 6 | DAPL1 | 3,351 |
| 7 | **LEF1** | 1,848 |
| 8 | KIF26A | 1,246 |
| 9 | TNFRSF19 | 1,148 |
| 10 | SLC9A2 | 702 |
| 11 | SGK3 | 363 |
| 12 | PCDH19 | 360 |
| 13 | ENSG00000243491 | 327 |
| 14 | SLCO5A1 | 285 |
| 15 | CALN1 | 172 |
| 16 | NALCN | 108 |
| 17 | ENSG00000286069 | 89 |
| 18 | BHLHE40-AS1 | 85 |
| 19 | ALDH8A1 | 70 |
| 20 | GLRA2 | 26 |

**Table 3:**
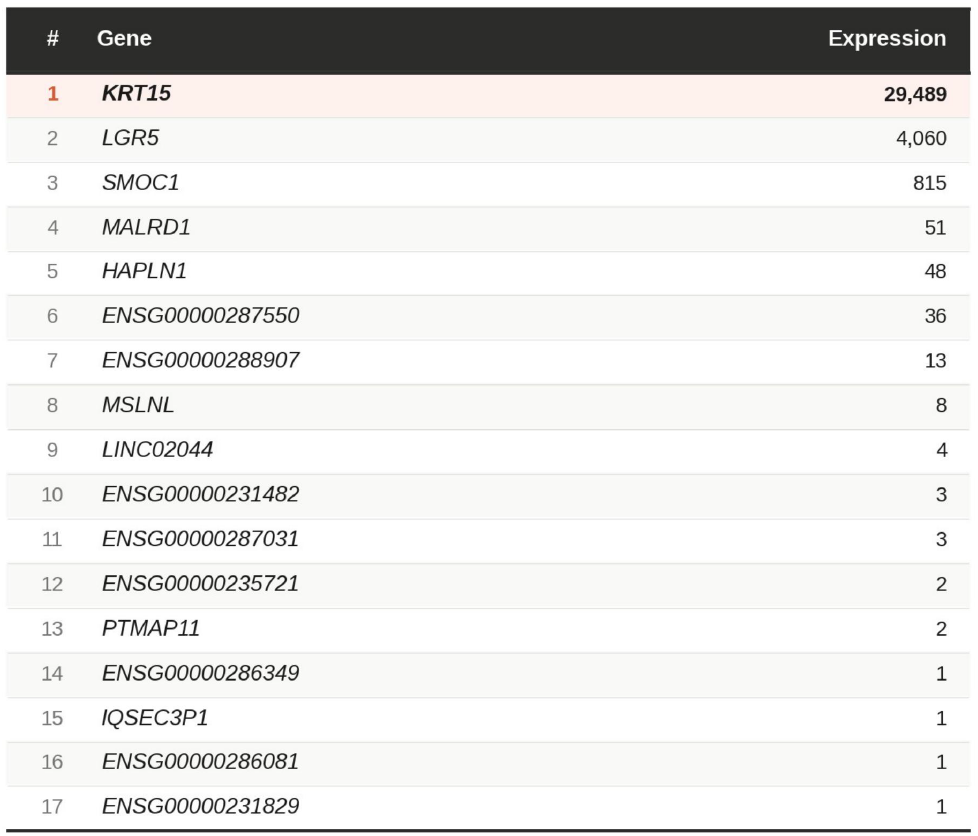
Randomised KRT15 spectral prompt. Output from a single randomly drawn spectral level. Expression shown as mean normalised read count.

**Table 4:**
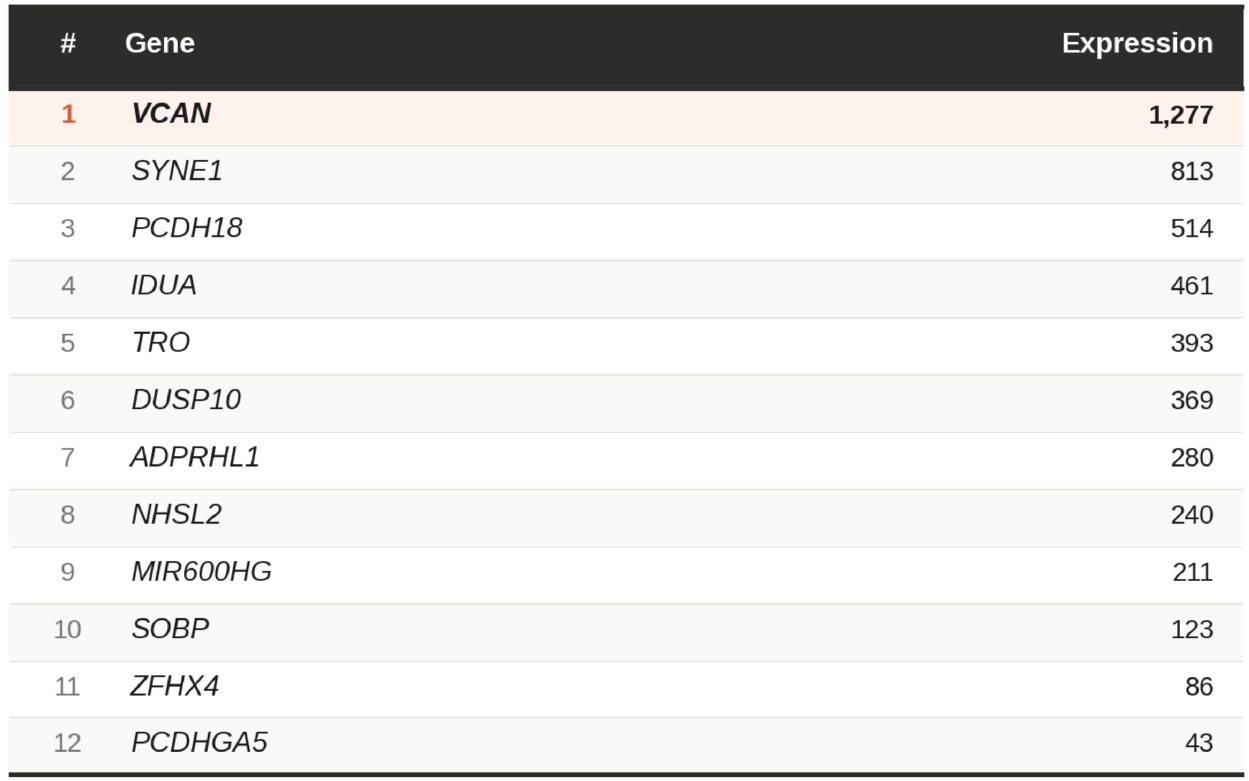
Randomised VCAN spectral prompt. Output from a single randomly drawn spectral level. Expression shown as mean normalised read count.

**Table 5:** Randomised DCT spectral prompt. Output from a single randomly drawn spectral level. Expression shown as mean normalised read count.

| # | Gene | Expression |
| --- | --- | --- |
| 1 | **DCT** | 540 |
| 2 | TMPRSS4 | 474 |
| 3 | TMC5 | 298 |
| 4 | SPRR2F | 237 |
| 5 | ACKR2 | 84 |
| 6 | DUSP9 | 42 |
| 7 | ENSG00000286299 | 24 |
| 8 | XCR1 | 15 |
| 9 | LINC02178 | 11 |
| 10 | FAM182A | 7 |
| 11 | ADARB2 | 6 |
| 12 | ENSG00000250762 | 6 |
| 13 | LINC02660 | 3 |
| 14 | NCF4-AS1 | 3 |

**Table 6:**
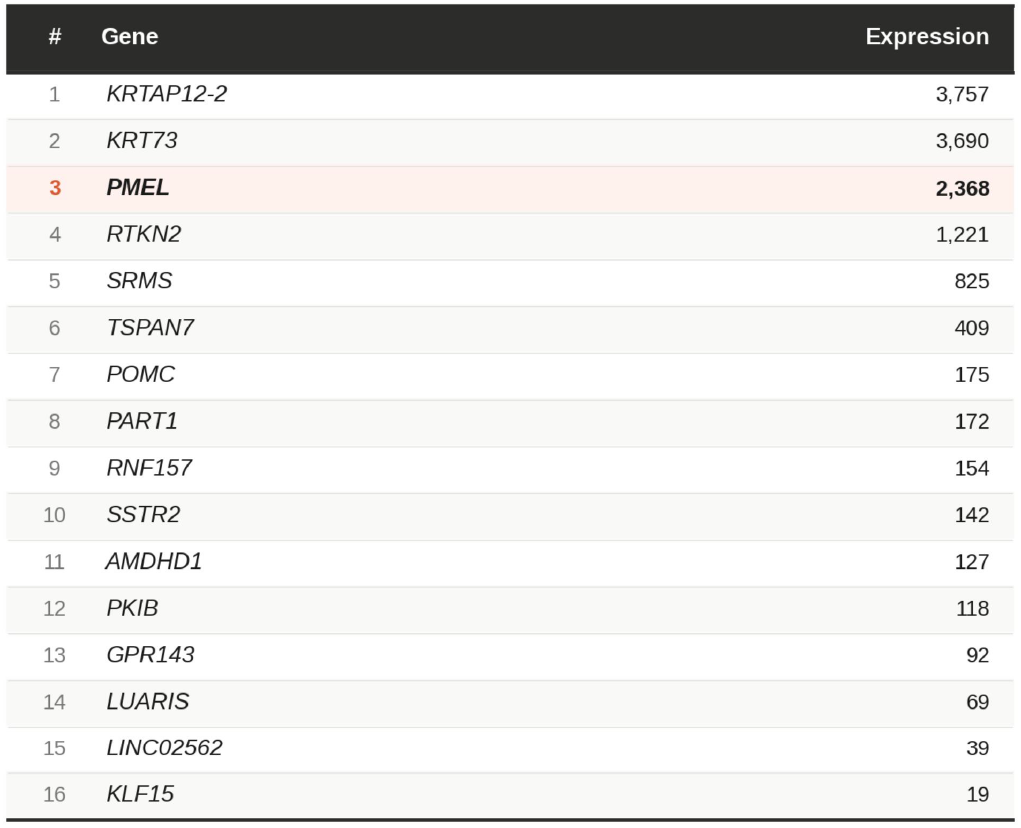
Randomised PMEL spectral prompt. Output from a single randomly drawn spectral level. Expression shown as mean normalised read count.

| # | Gene | Expression |
| --- | --- | --- |
| 1 | KRTAP12-2 | 3,757 |
| 2 | KRT73 | 3,690 |
| 3 | **PMEL** | 2,368 |
| 4 | RTKN2 | 1,221 |
| 5 | SRMS | 825 |
| 6 | TSPAN7 | 409 |
| 7 | POMC | 175 |
| 8 | PART1 | 172 |
| 9 | RNF157 | 154 |
| 10 | SSTR2 | 142 |
| 11 | AMDHD1 | 127 |
| 12 | PKIB | 118 |
| 13 | GPR143 | 92 |
| 14 | LUARIS | 69 |
| 15 | LINC02562 | 39 |
| 16 | KLF15 | 19 |

**Table 7:** Randomised MKI67 spectral prompt. Output from a single randomly drawn spectral level. Expression shown as mean normalised read count.

| # | Gene | Expression |
| --- | --- | --- |
| 1 | **MKI67** | **4,039** |
| 2 | TOP2A | 3,029 |
| 3 | ATAD2 | 1,679 |
| 4 | LMNB1 | 1,640 |
| 5 | CCNA2 | 1,120 |
| 6 | KIF11 | 1,118 |
| 7 | TMEM97 | 1,096 |
| 8 | PRR11 | 952 |
| 9 | BRCA1 | 931 |
| 10 | CEP55 | 848 |
| 11 | E2F2 | 785 |
| 12 | SHCBP1 | 736 |
| 13 | CENPE | 673 |

**Table 8:**
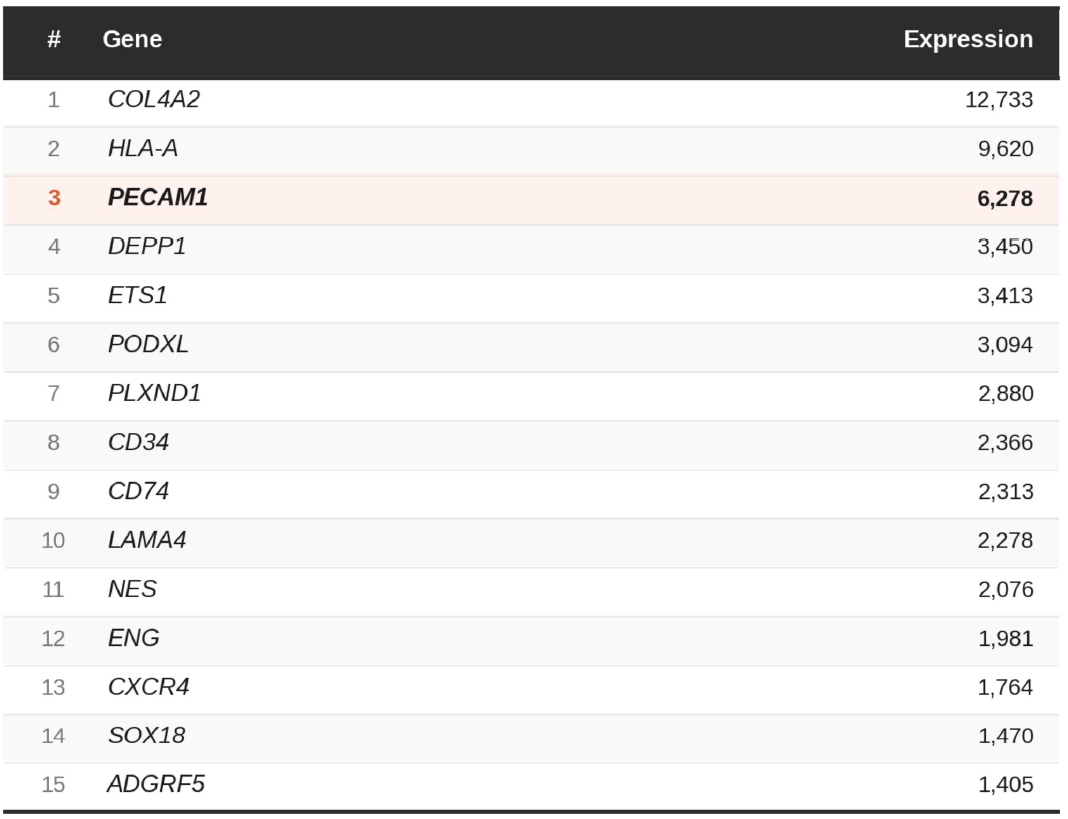
Randomised PECAM1 spectral prompt. Output from a single randomly drawn spectral level. Expression shown as mean normalised read count.

**Table 9:**
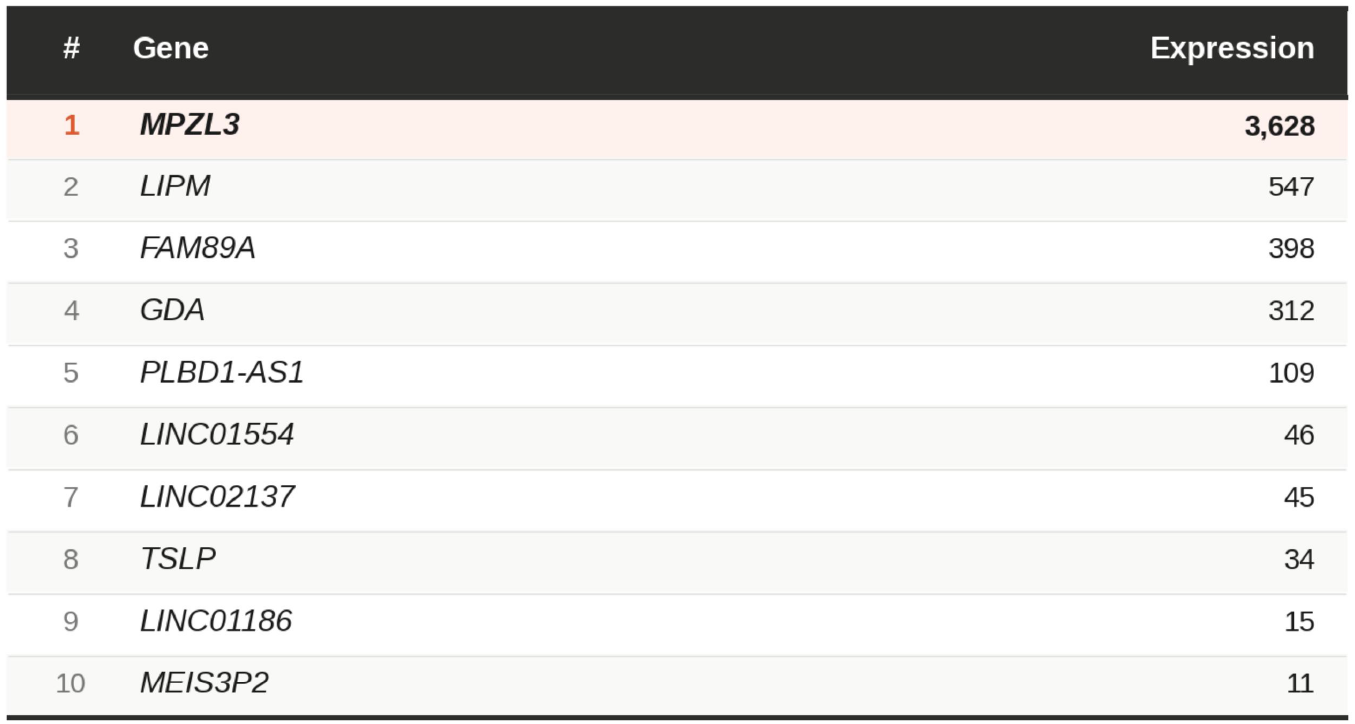
Randomised MPZL3 spectral prompt. Output from a single randomly drawn spectral level. Expression shown as mean normalised read count.

| # | Gene | Expression |
| --- | --- | --- |
| 1 | **MPZL3** | **3,628** |
| 2 | LIPM | 547 |
| 3 | FAM89A | 398 |
| 4 | GDA | 312 |
| 5 | PLBD1-AS1 | 109 |
| 6 | LINC01554 | 46 |
| 7 | LINC02137 | 45 |
| 8 | TSLP | 34 |
| 9 | LINC01186 | 15 |
| 10 | MEIS3P2 | 11 |

KRT85 returned KRT35 and LEF1 (Table 1), whereas LEF1 reciprocally returned KRT85 and other matching genes e.g. LAP3 and SP6, as well as uniquely returning DSG4 for this spectral prompt (Table 2). For the remaining prompts, examples of returned results were: LGR5, SMOC1 (KRT15; Table 3); NHSL2, PCDH18 (VCAN; Table 4); SPRR2F, TMPRSS4 (DCT; Table 5); POMC, GPR143 (PMEL; Table 6); TOP2A, BRCA1 (MKI67; Table 7); ETS1, CD34 (PECAM1; Table 8); FAM89A, TSLP (MPZL3; Table 9).

Subsequently for each marker I swept all non-degenerative levels of two constructed manifolds (varied by ARPACK seed). Returned genes were aggregated and expressed as a percentage for recurrence. The top 7 recurrent genes per prompt were plotted (minimum read count 10 shown to demonstrate potential emergent relationships not dominated by expression level) (Figures 1–9), and the remaining returns (all invariant between the two manifold constructions), were listed in Tables 10–18.

**Table 10:**
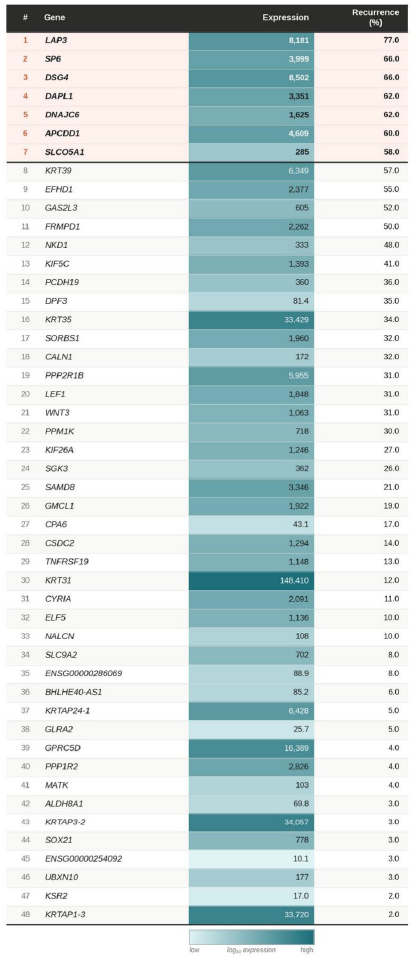
KRT85 aggregated spectral prompt. All genes invariant across both manifold constructions. Bold: top 7 by recurrence. Expression coloured by log_10_ scale.

**Table 11:**
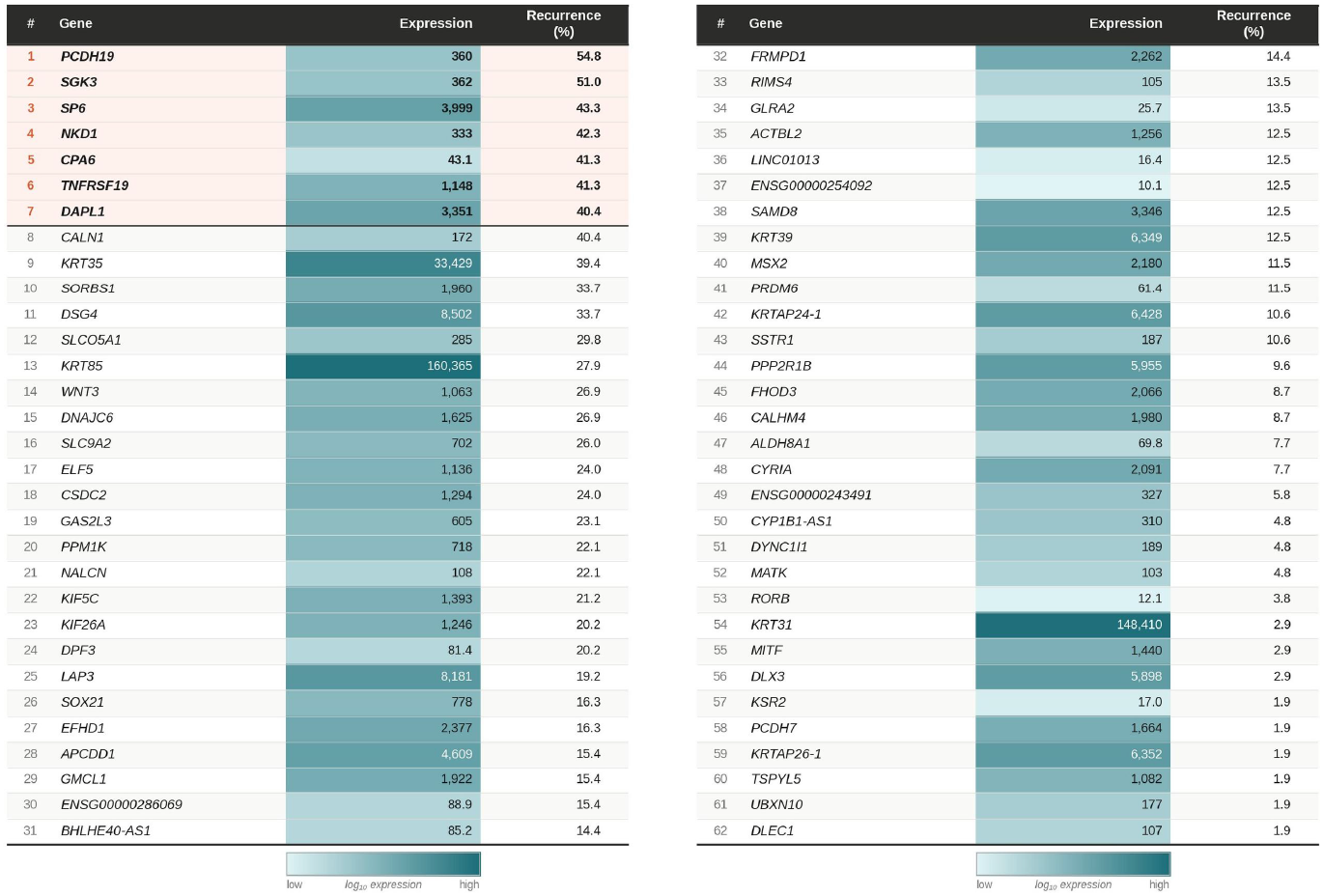
LEF1 aggregated spectral prompt. All genes invariant across both manifold constructions. Bold: top 7 by recurrence. Expression coloured by log_10_ scale.

**Table 12:**
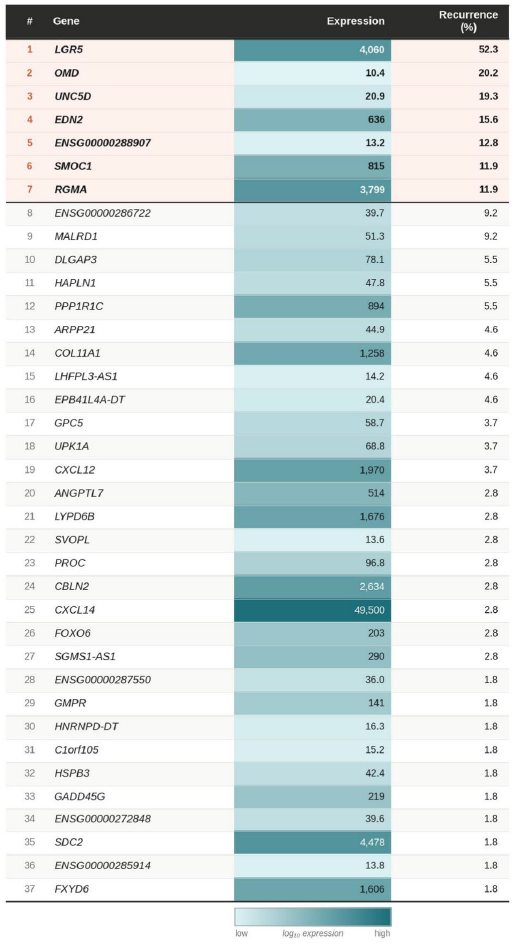
KRT15 aggregated spectral prompt. All genes invariant across both manifold constructions. Bold: top 7 by recurrence. Expression coloured by log_10_ scale.

**Table 13:**
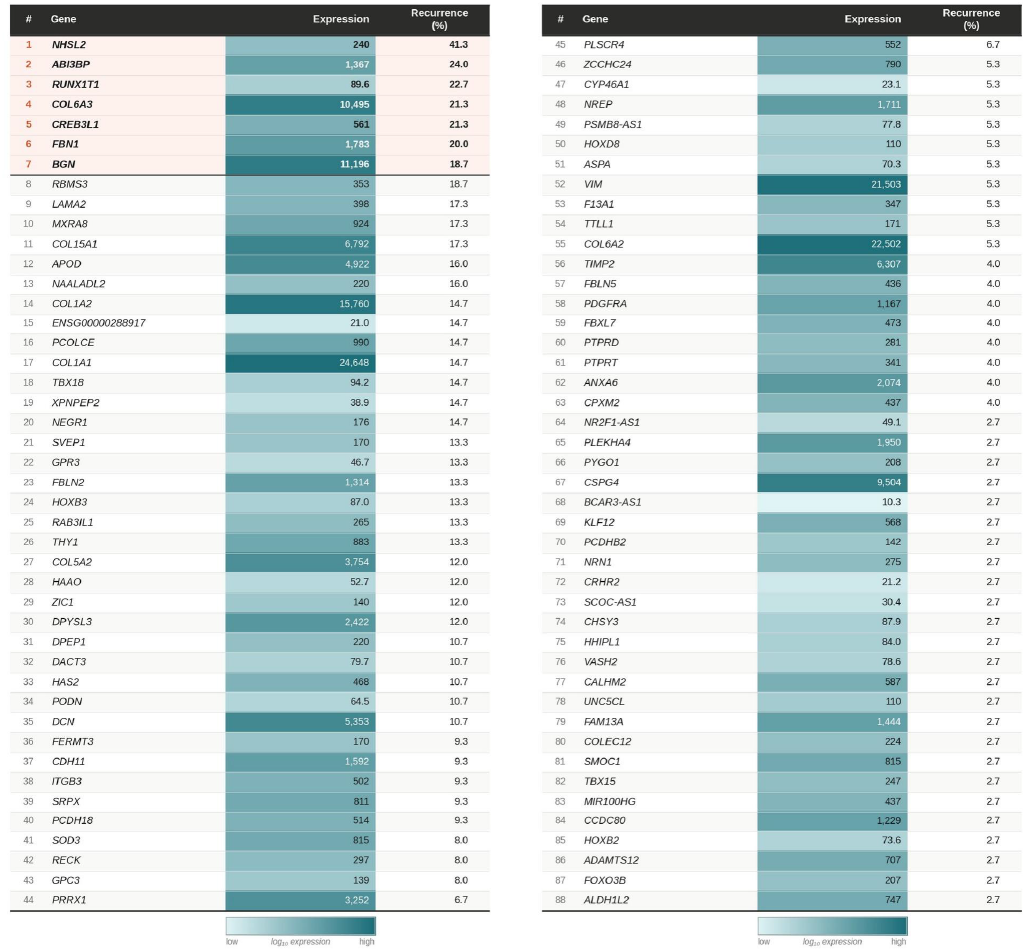
VCAN aggregated spectral prompt. All genes invariant across both manifold constructions. Bold: top 7 by recurrence. Expression coloured by log_10_ scale.

**Table 14:**
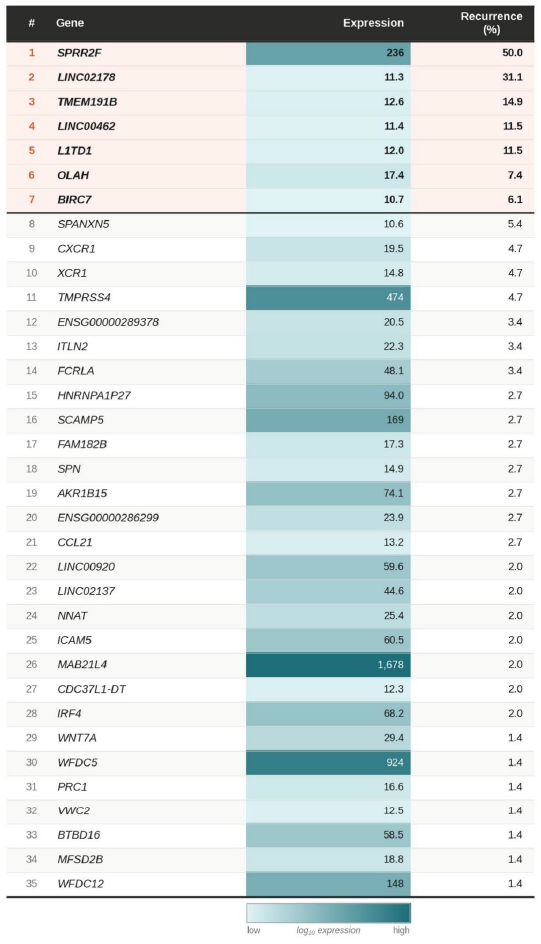
DCT aggregated spectral prompt. All genes invariant across both manifold constructions. Bold: top 7 by recurrence. Expression coloured by log_10_ scale.

**Table 15:**
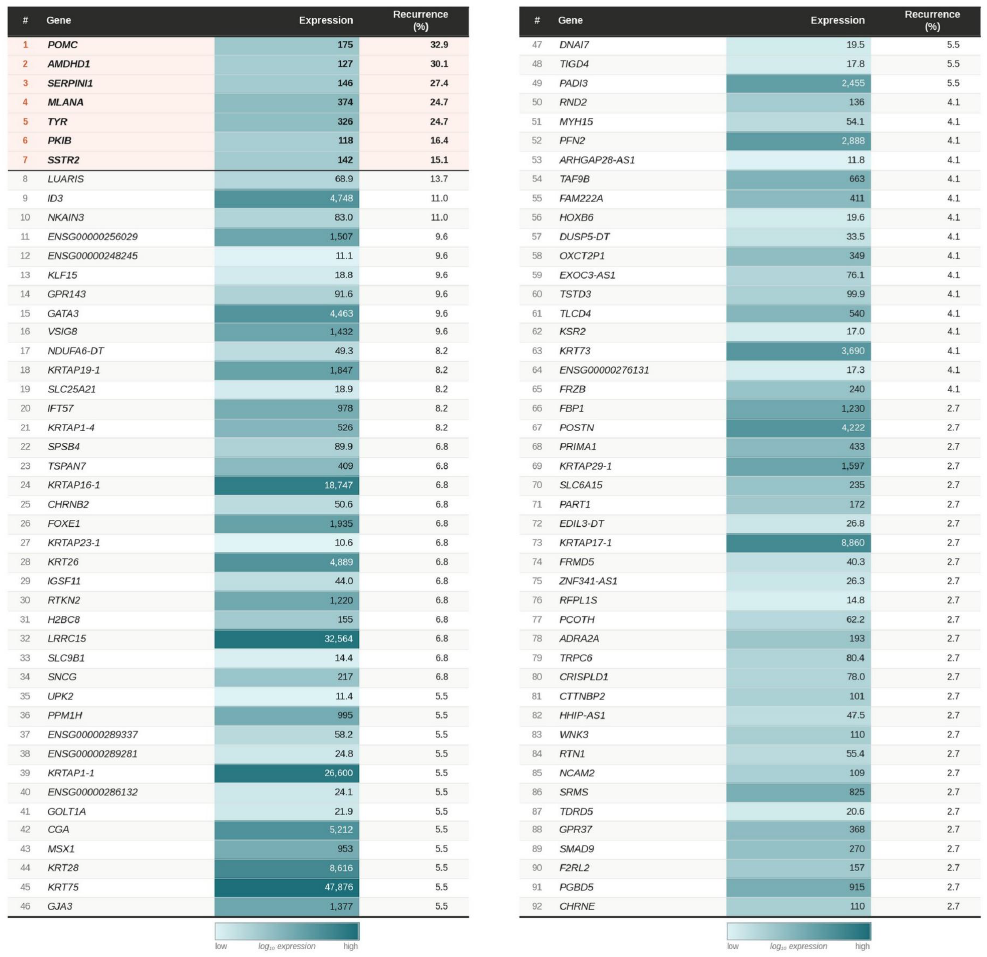
PMEL aggregated spectral prompt. All genes invariant across both manifold constructions. Bold: top 7 by recurrence. Expression coloured by log_10_ scale.

**Table 16:**
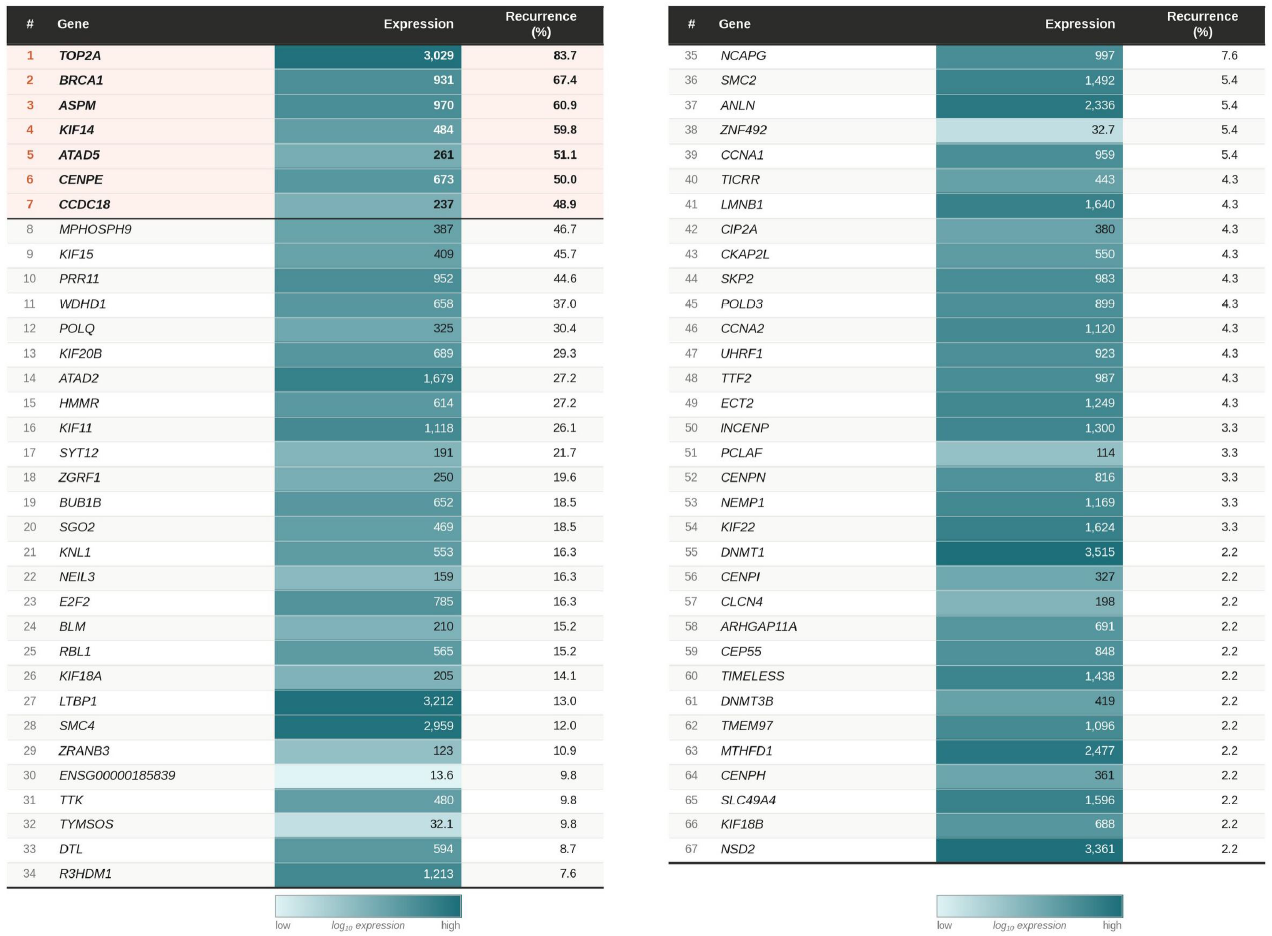
MKI67 aggregated spectral prompt. All genes invariant across both manifold constructions. Bold: top 7 by recurrence. Expression coloured by log_10_ scale.

**Table 17:**
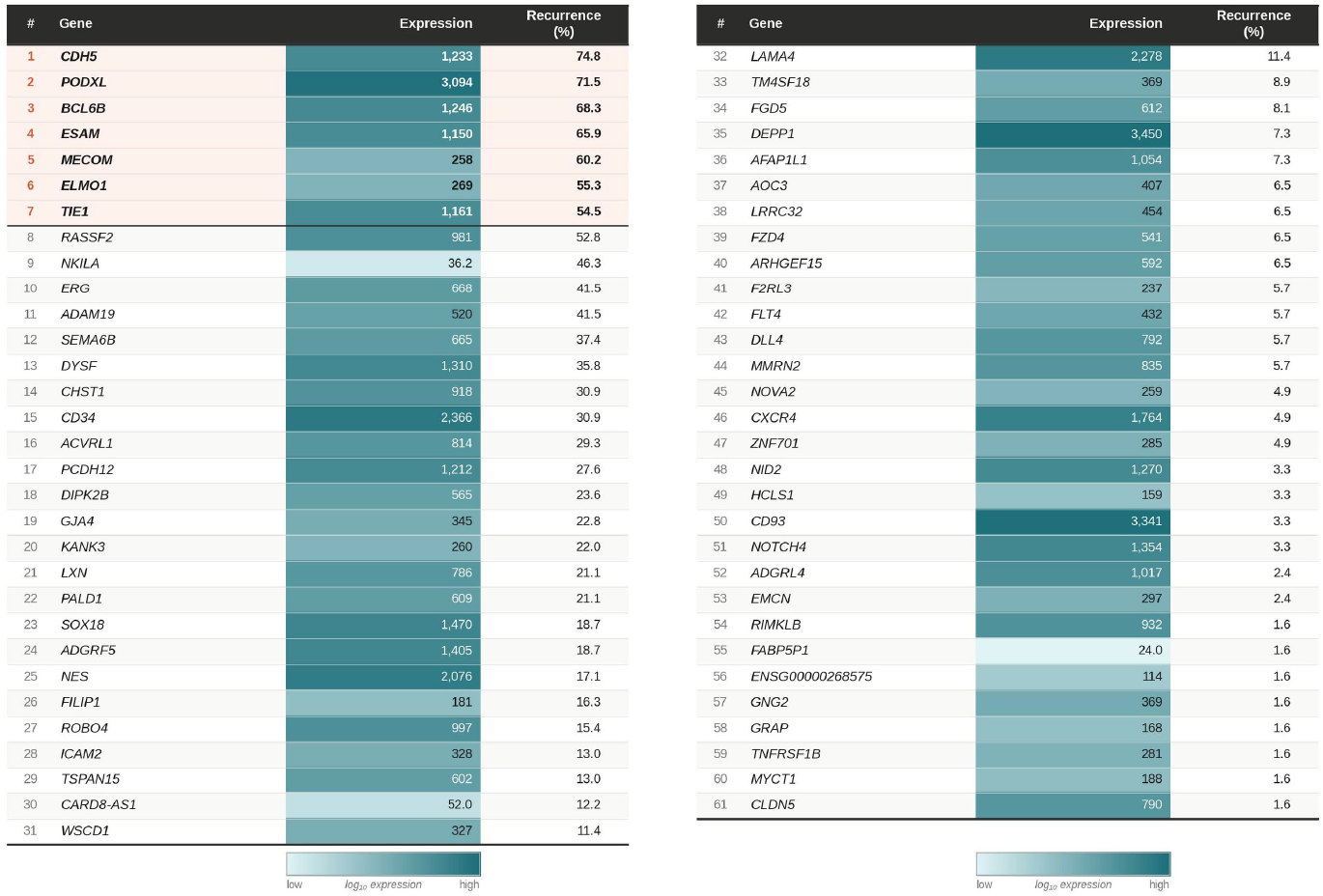
PECAM1 aggregated spectral prompt. All genes invariant across both manifold constructions. Bold: top 7 by recurrence. Expression coloured by log_10_ scale.

**Table 18:**
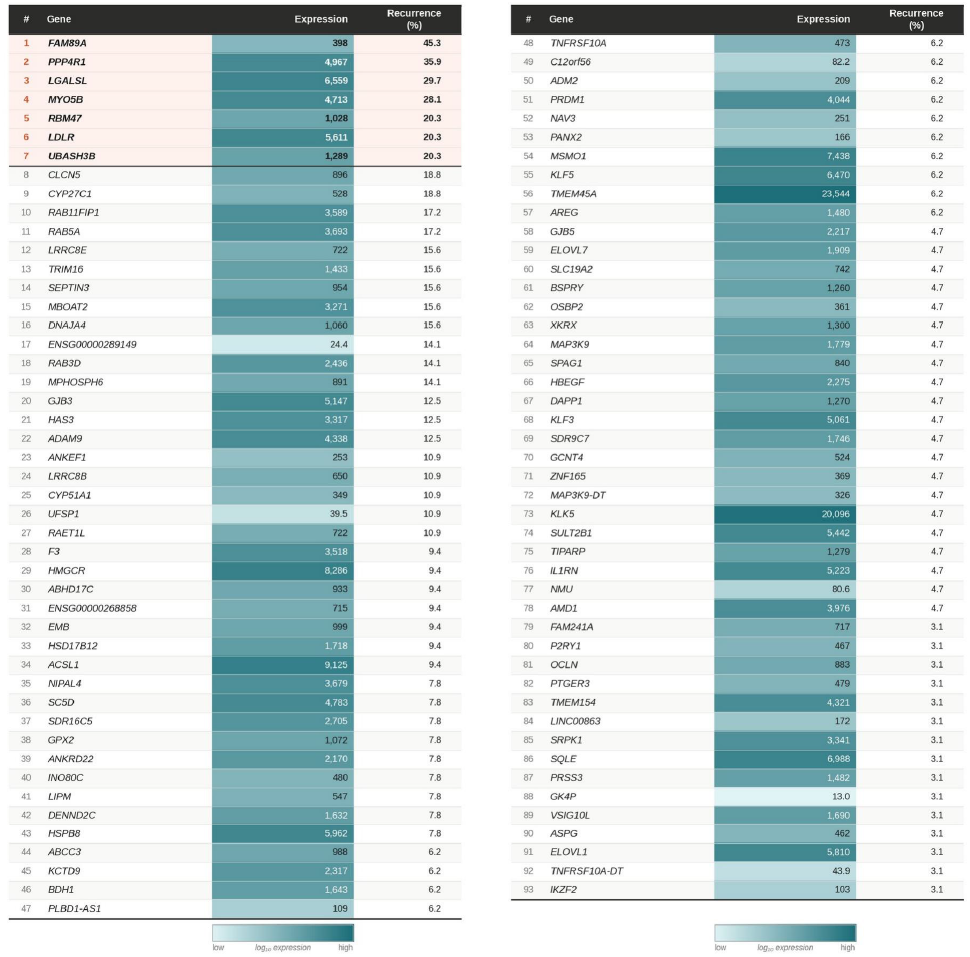
MPZL3 aggregated spectral prompt. All genes invariant across both manifold constructions. Bold: top 7 by recurrence. Expression coloured by log_10_ scale.

**Figure 1.**
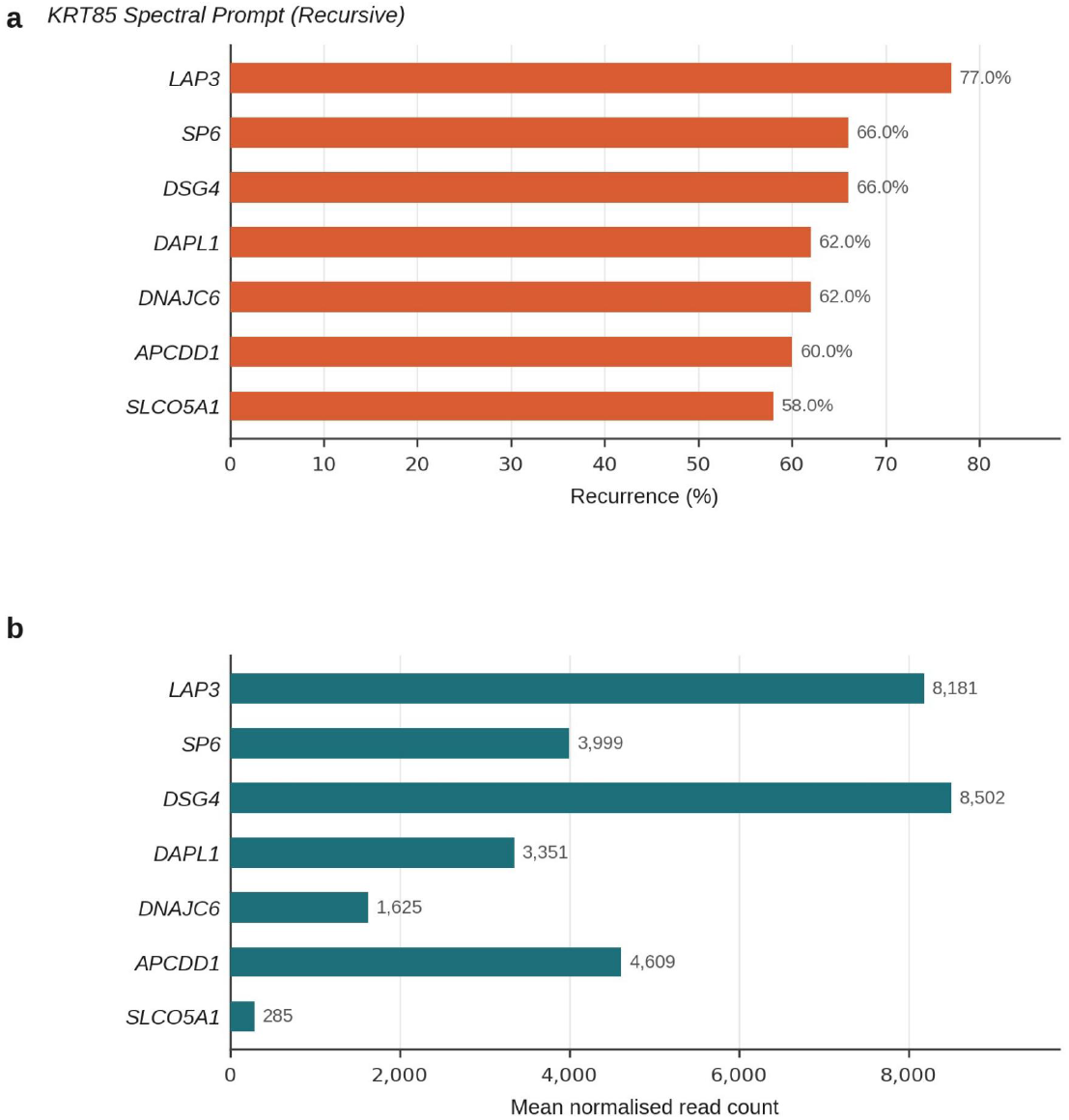
KRT85 aggregated spectral prompt. Top 7 recurrent genes by percentage across two manifold constructions. (a) Recurrence. (b) Mean normalised read count.

**Figure 2.**
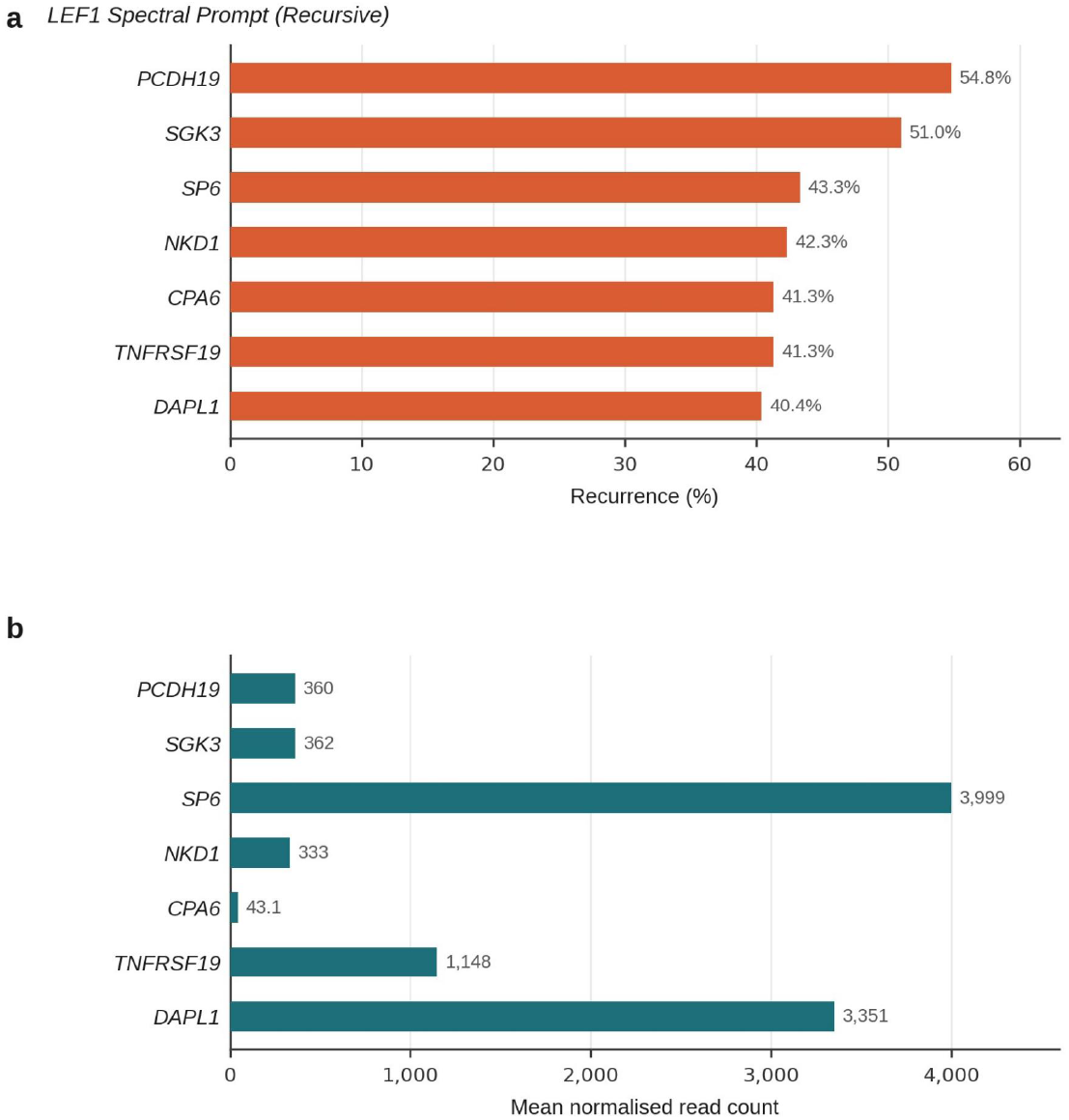
LEF1 aggregated spectral prompt. Top 7 recurrent genes by percentage across two manifold constructions. (a) Recurrence. (b) Mean normalised read count.

**Figure 3.**
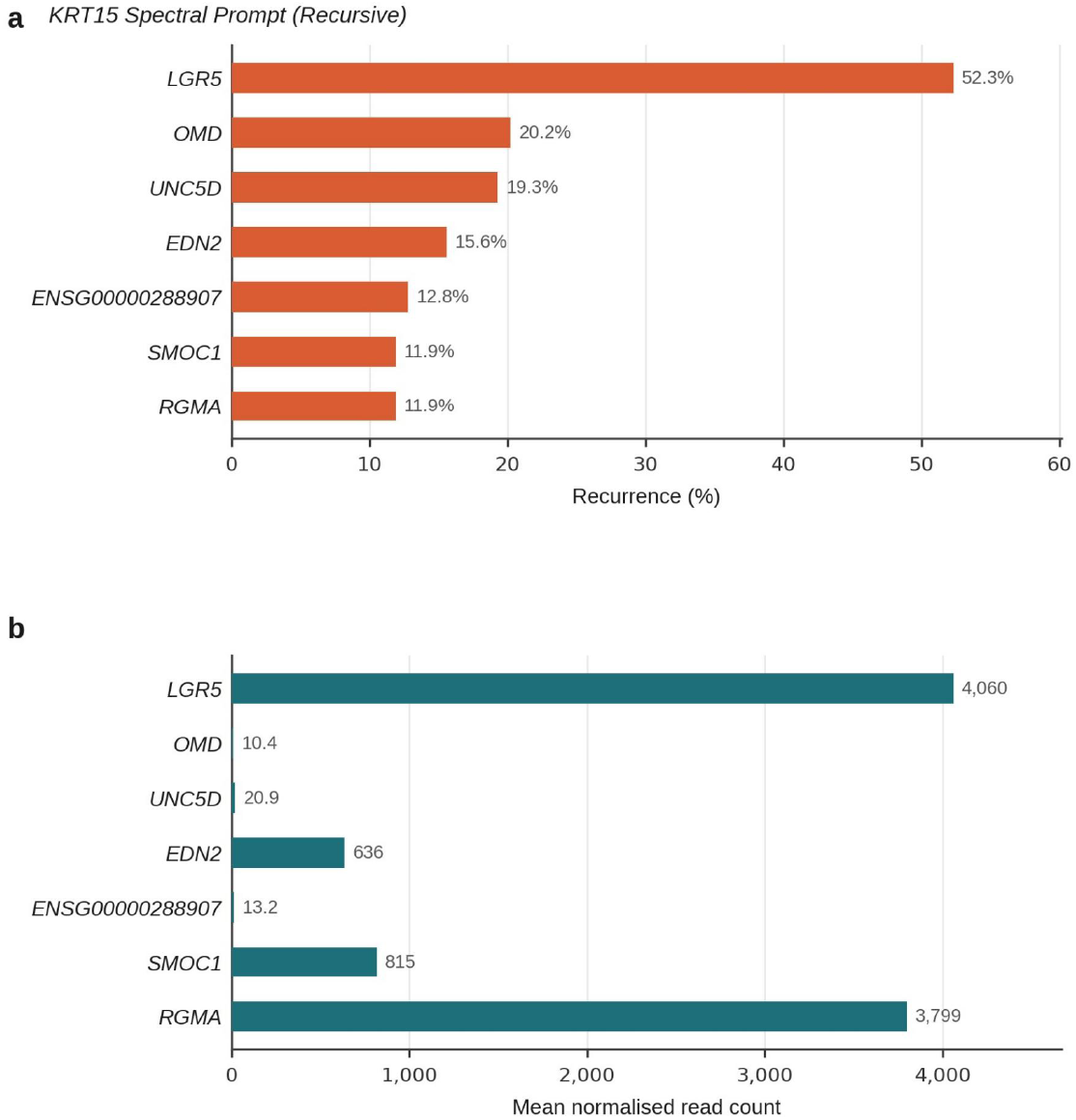
KRT15 aggregated spectral prompt. Top 7 recurrent genes by percentage across two manifold constructions. (a) Recurrence. (b) Mean normalised read count.

**Figure 4.**
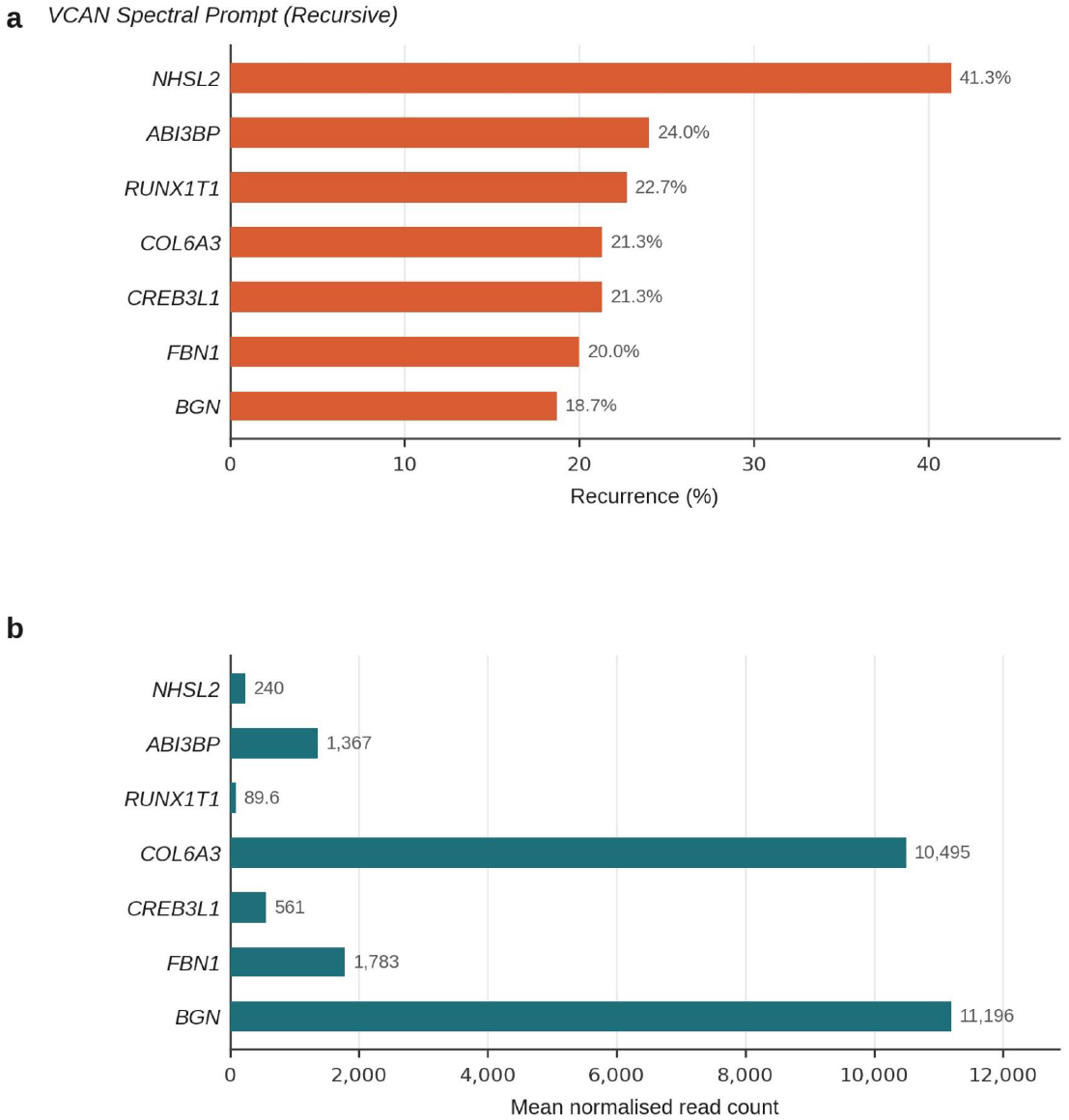
VCAN aggregated spectral prompt. Top 7 recurrent genes by percentage across two manifold constructions. (a) Recurrence. (b) Mean normalised read count.

**Figure 5.**
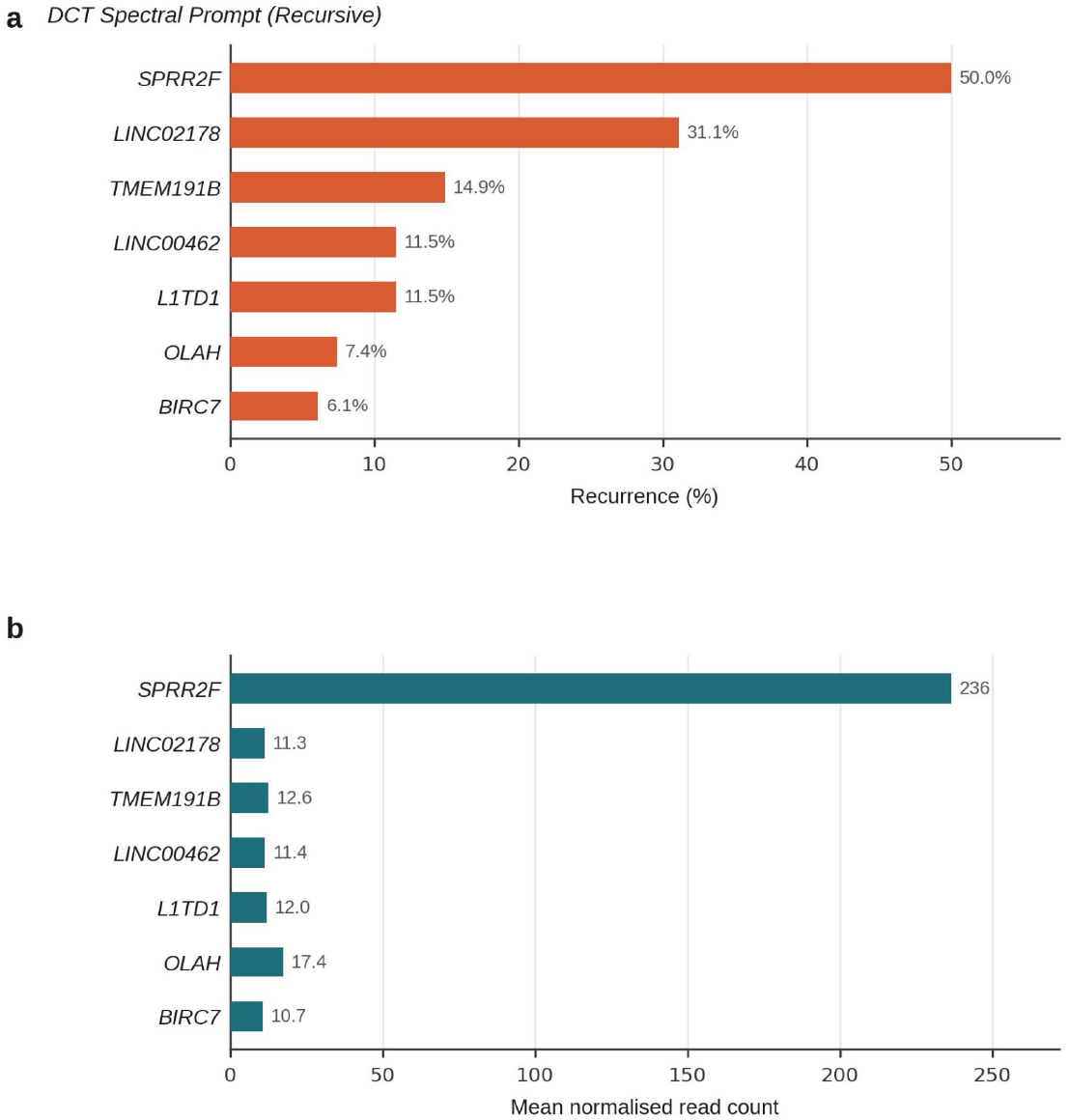
DCT aggregated spectral prompt. Top 7 recurrent genes by percentage across two manifold constructions. (a) Recurrence. (b) Mean normalised read count.

**Figure 6.**
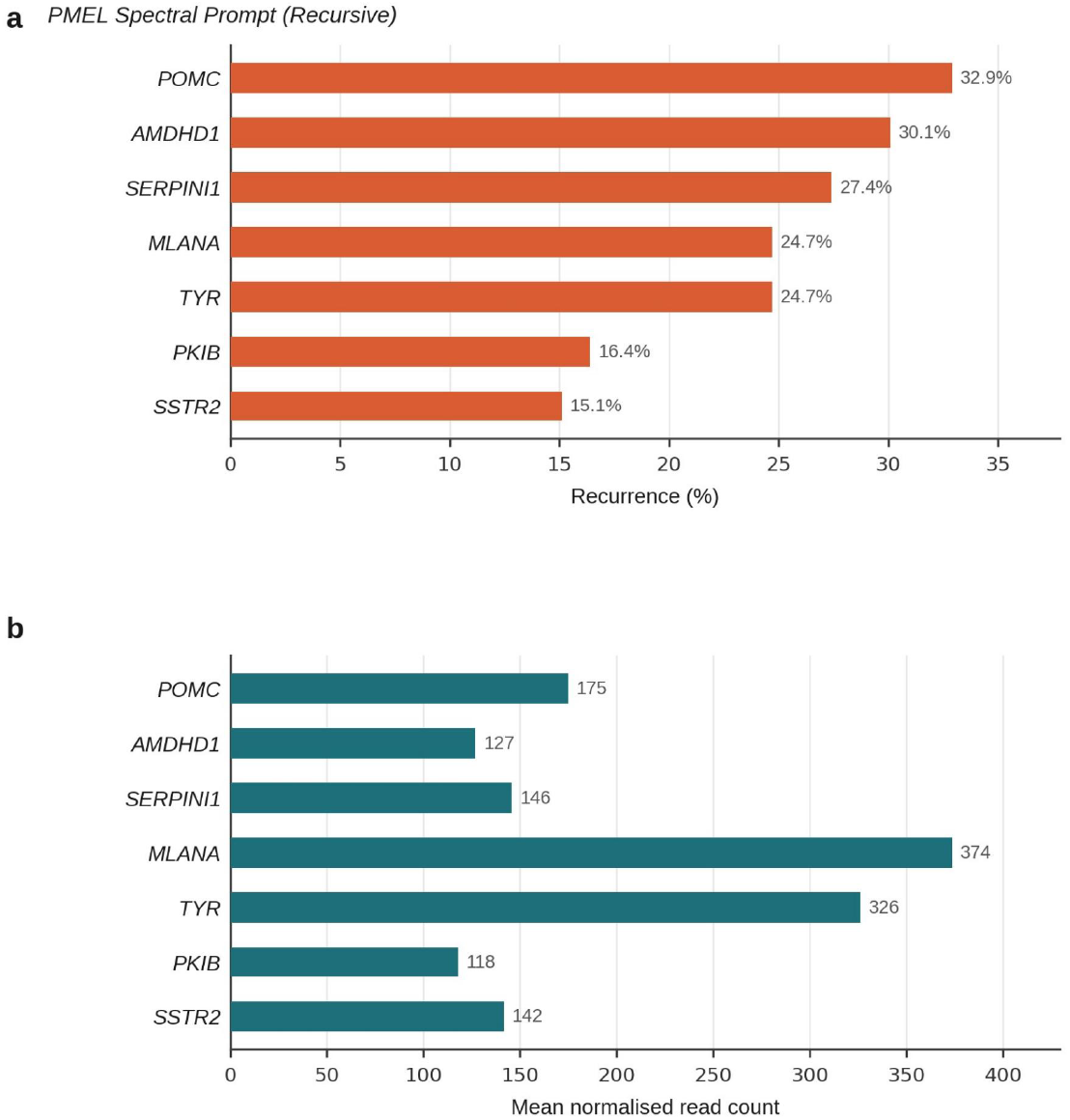
PMEL aggregated spectral prompt. Top 7 recurrent genes by percentage across two manifold constructions. (a) Recurrence. (b) Mean normalised read count.

**Figure 7.**
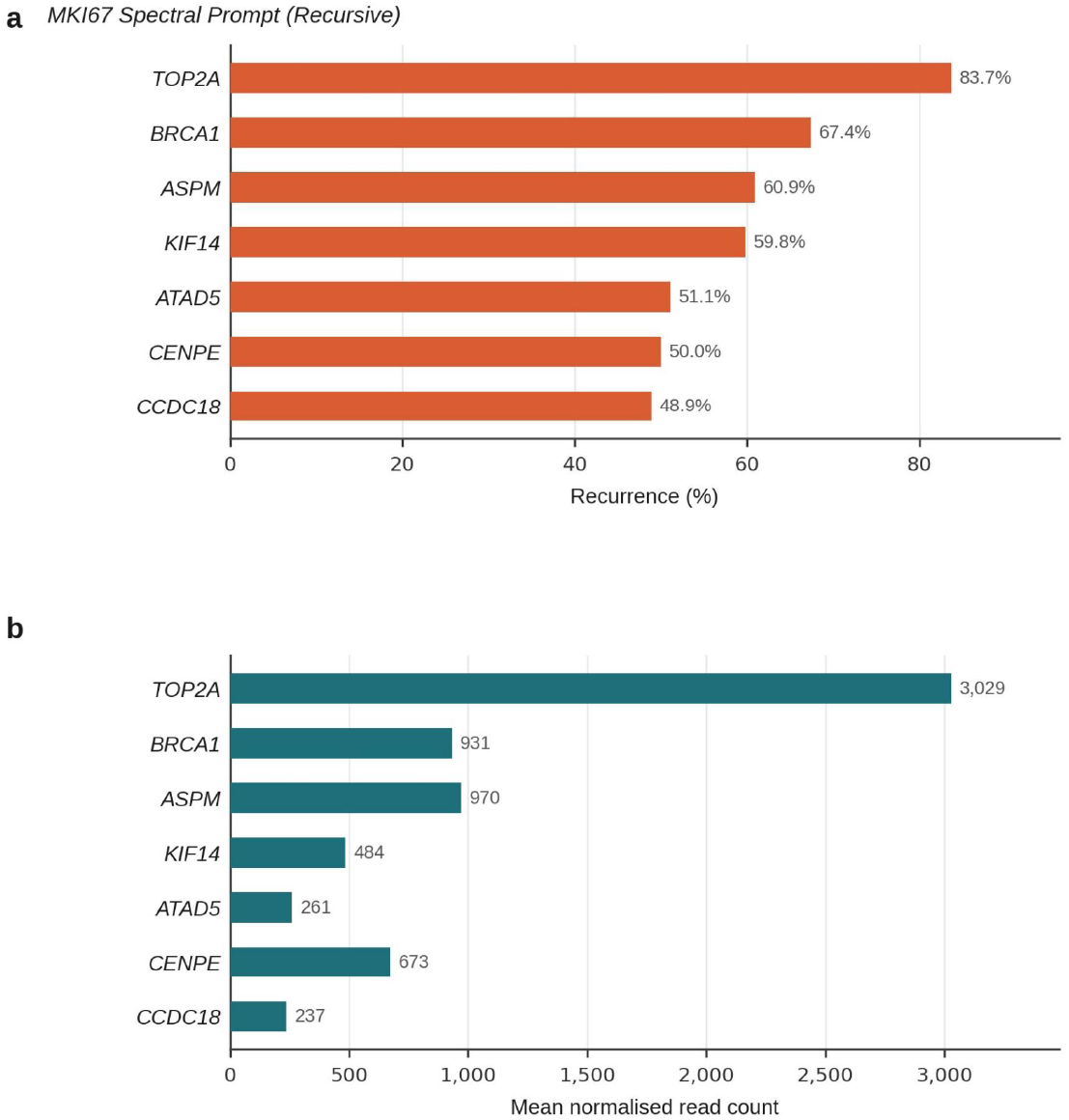
MKI67 aggregated spectral prompt. Top 7 recurrent genes by percentage across two manifold constructions. (a) Recurrence. (b) Mean normalised read count.

**Figure 8.**
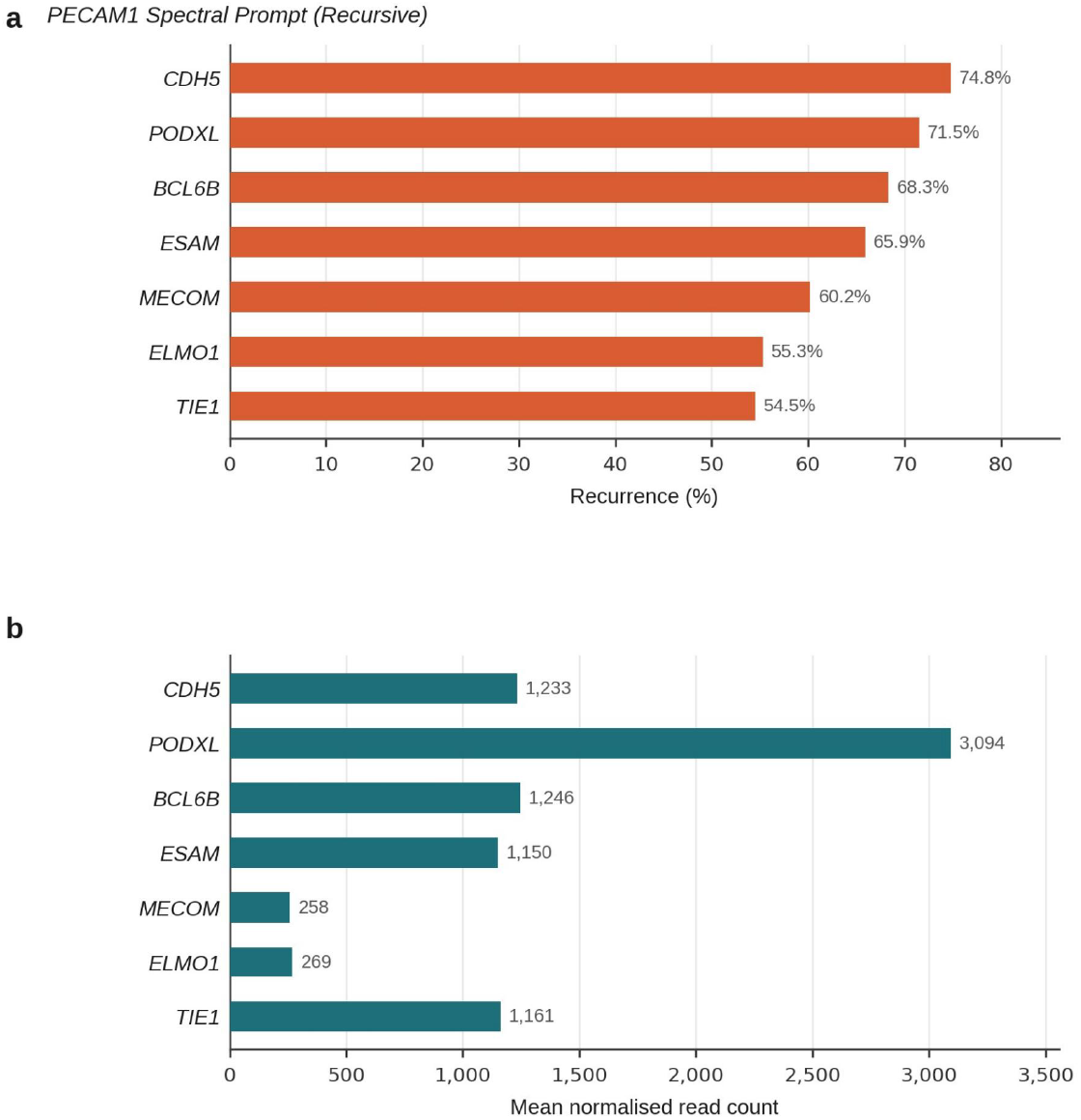
PECAM1 aggregated spectral prompt. Top 7 recurrent genes by percentage across two manifold constructions. (a) Recurrence. (b) Mean normalised read count.

**Figure 9.**
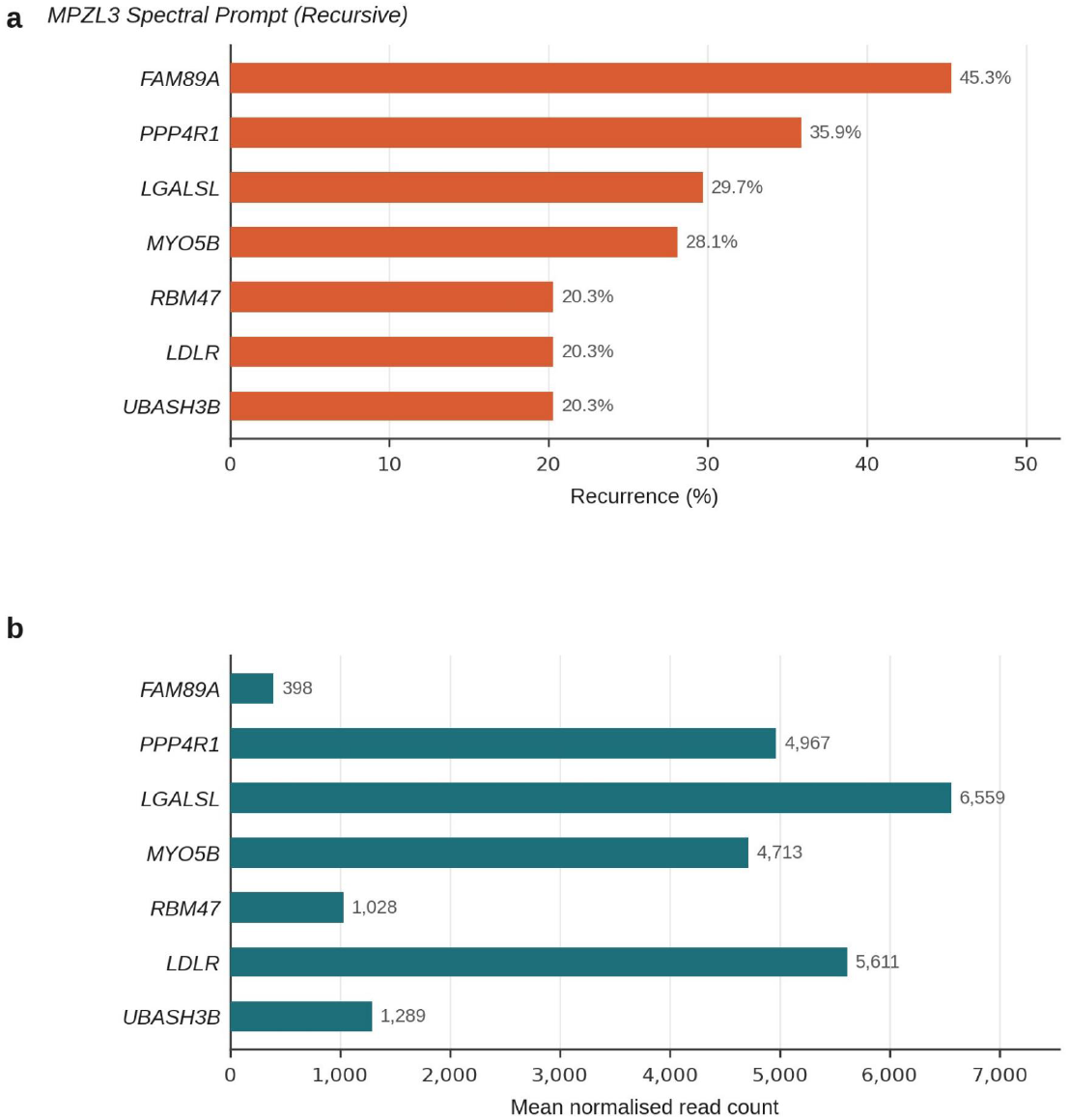
MPZL3 aggregated spectral prompt. Top 7 recurrent genes by percentage across two manifold constructions. (a) Recurrence. (b) Mean normalised read count.

Following aggregation, KRT85 returned APCDD1, implicated in hereditary hypotrichosis simplex and expressed in human hair follicles [19], and DAPL1, previously predicted to be expressed in the hair shaft cortex by single-cell profiling of the human hair follicle [20] (Figure 1, Table 10). LEF1 returned NKD1 and SGK3, Wnt/*β*-catenin signalling-associated genes [21, 22] (Figure 2, Table 11). The top aggregated hits for PECAM1 and MKI67 produced clear endothelial and mitosis signatures (Figures 8, 7), VCAN aggregated top hits included known ECM genes COL6A3, FBN1, BGN (Figure 4). For MPZL3, the top hits produced an as yet unclear collective profile; however, by expanding the cutoff to *>*10% recurrence, a Rab-associated endosomal recycling/exocytic signature can be seen (MYO5B, RAB5A, RAB11FIP1, RAB3D) (Table 18).

DCT produced top hits that were not immediately legible to author’s intended use as a melanocyte biology gene marker in human hair follicles. However, a single DCT prompt did return IRF4, a MITF target gene [23] (Table 14). When PMEL was alternatively employed, this produced melanocyte and melanogenesis associated genes (MLANA, TYR, POMC) (Figure 6, Table 15).

The aggregated signature for KRT15 cemented LGR5 as a top recurrent gene (found in 52.3% of outputs), alongside EDN2 [24], SMOC1, and RGMA, which recurred ∼11–15% (omitting low expression genes) (Figure 3, Table 12). Prior to the development of the aggregated method to capture spectral outputs from gene prompts across modes in the manifold, individual prompts returned lists that included CALML3 and MUCL1 in one output (Table 19), and CXCL14, RGMA and LHX2 in another output (Table 20) (where LHX2 and MUCL1 were found to be not invariant to *>*1 manifold constructions).

**Table 19:** Individual KRT15 spectral prompt output (pre-aggregation).

| # | Gene | Expression |
| --- | --- | --- |
| 1 | *KRT15* | 29,489 |
| 2 | *CALML3* | 71,052 |
| 3 | *MUCL1* | 25,112 |
| 4 | *TUBA1A* | 14,445 |
| 5 | *MGP* | 6,837 |
| 6 | *SDC2* | 4,478 |
| 7 | *EFEMP1* | 3,977 |
| 8 | *CBLN2* | 2,634 |
| 9 | *CXCL12* | 1,971 |
| 10 | *LYPD6B* | 1,676 |

**Table 20:** Individual KRT15 spectral prompt output (pre-aggregation).

| # | Gene | Expression |
| --- | --- | --- |
| 1 | *KRT15* | 29,489 |
| 2 | *CXCL14* | 49,500 |
| 3 | *RGMA* | 3,799 |
| 4 | *LHX2* | 3,407 |
| 5 | *CTNND2* | 1,564 |
| 6 | *PPP1R1C* | 894 |
| 7 | *FGF18* | 714 |
| 8 | *CNTN4* | 325 |
| 9 | *GADD45G* | 219 |
| 10 | *FOXO6* | 203 |
| 11 | *ENSG00000285914* | 14 |
| 12 | *SVOPL* | 14 |

In previous work, we found that LHX2 was expressed sporadically across the human hair follicle ORS [25] and is known in eHFSC biology [26], suggesting that this variation does not preclude the detection of real biological relationships, and may represent weaker statistical association versus identified recurrent/invariant signatures. Indeed, despite not surviving *>*1 manifold construction, *in situ* immunofluorescent tissue section staining of MUCL1 in human hair follicle tissue sections showed that it localised to the ORS, including in some instances within KRT15+ cells, and/or below the bulge compartment (Figure 10). Motivated by the outputs of these early results (pre-aggregation), dual staining for KRT15 alongside CXCL14 or RGMA was conducted (Figure 10). This confirmed human hair follicle anagen bulge expression of CXCL14 [20] and newly identified the presence of RGMA protein expression (involved with BMP and SMAD1/5/8 signalling [27, 28]) within KRT15+ cells, which was seen in some follicles to concentrate to the upper portion of the KRT15+ bulge region.

**Figure 10.**
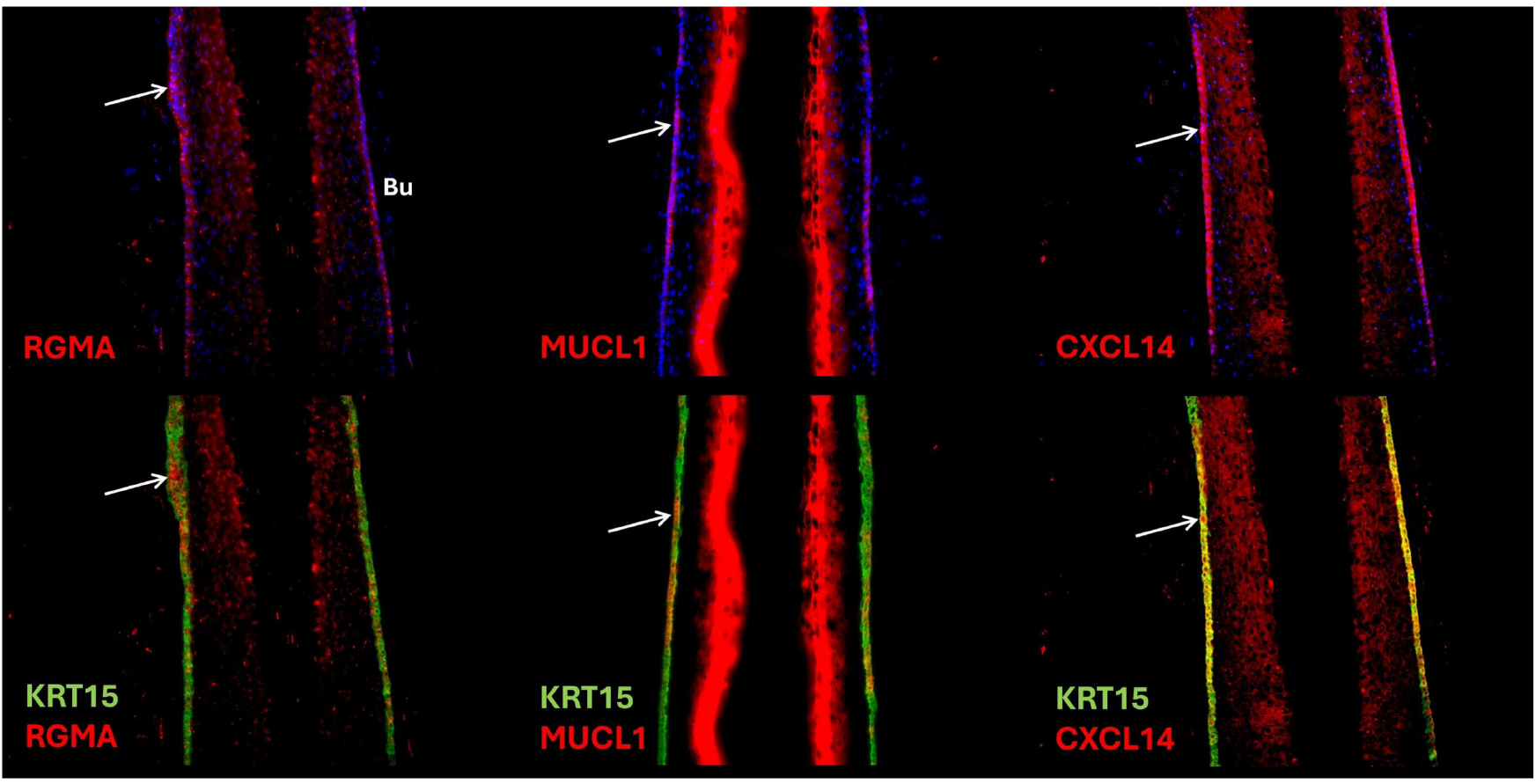
*In situ* human hair follicle bulge immunofluorescence for RGMA, MUCL1 and CXCL14. Top: single channel with DAPI. Bottom: dual stain with KRT15. Arrows indicate co-localising staining patterns. Bu, bulge.

### Spectral Prompting Enhances Single-Cell Analyses Without Clustering

As a demonstration that this approach is effective on single-cell data, I used the same pipeline to construct a manifold from data obtained from GSE129611 [20], specifically the 10x sample HFU13. I then deployed the same Spectral Prompting system (independent from the use of clustering, dimensionality reduction or reference atlases), using KRT15 as a seed on this single manifold. As an illustrative example, one individual prompt produced (alongside KRT15 itself) the genes CYP1B1, FHL2, PPP1R1C, LMCD1, CHST2, FAM13A, CXCL14, DDAH2, ITM2A, SFRP1, LHX2, NET1, FXYD6, DIO2, HCFC1R1, CBX1, MRPS6, RCAN1. From this result alone, several markers here were notable to the author, namely CXCL14 and LHX2 as discussed above, but also DIO2, identified in a seminal human eHFSC study by Ohyama et al. [29] and the Wnt inhibitor SFRP1, studied in human hair follicles by Hawkshaw et al. [30].

Next I mapped this prompt return to individual cells, identifying 31 cells in this dataset that expressed all these markers (Figure 11), which were subsequently found to be enriched (against baseline expression by to 12.2x in *>*50% of cells) for BEX5, TGFB2, WIF1, ADAMTSL4, COL1A2, TUBB2B, CD200, EFEMP1, FBLN2, NPNT, SERPINE2, TNC and NFATC1.

**Figure 11.**
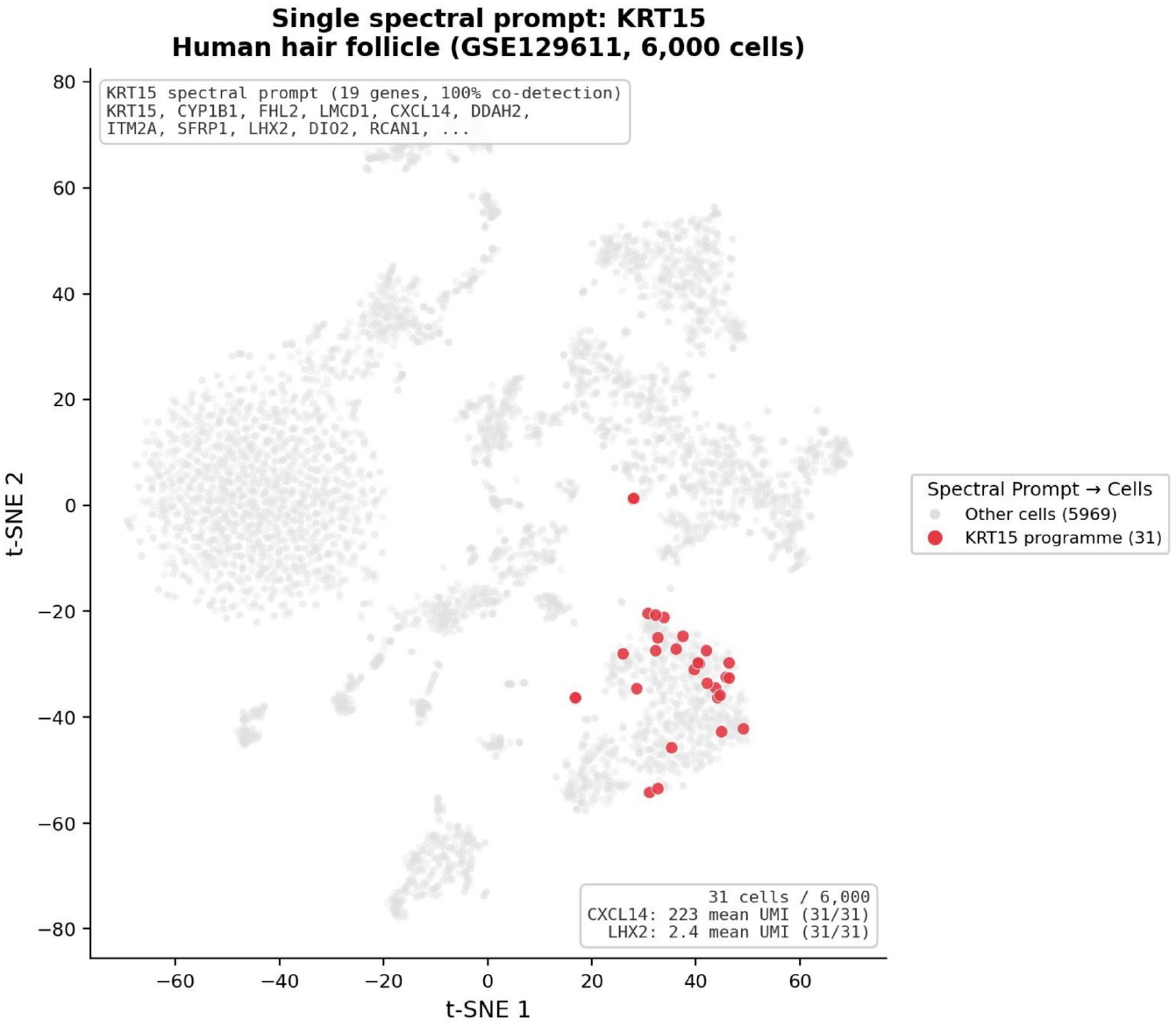
Single spectral prompt using KRT15 on single-cell data (GSE129611, 6,000 cells). 31 cells identified co-expressing all 19 returned genes.

### Technical note

The Spectral Prompting system can draw associations between transcripts that share similar relational structure on the manifold but lack functional relationship. Outputs should be independently validated. The pipeline described here has been implemented as a queryable discovery instrument (https://aethereos.net), Aethereos Biology, which operationalises these methods into distinct query modes across purpose-built data atlases. Moreover, whilst the technique can be deployed on relatively minimal data inputs (compared to foundation models), richer and more diverse datasets (i.e. from several patients with appropriate technical replicates) will produce richer atlases that carry greater confidence.

## Acknowledgments

This work and the Aethereos Biology system were independently developed by the author and received no dedicated project funding. Post-hoc *in situ* human hair follicle validation staining was kindly supported by the NIHR Manchester Biomedical Research Centre, and Derek Pye is thanked for performing these immunofluorescence experiments. The Genomic Technologies Core Facility, University of Manchester, is thanked for performing RNA-Seq experiments. Individual consenting patient donors, Dr Asim Shahmalak (The Crown Clinic, Manchester) and Dr Nilofer Farjo and Dr Bessam Farjo (Farjo Institute, Manchester) are thanked for providing tissue samples for research purposes.

